# Activation-induced deaminase is critical for the establishment of DNA methylation patterns prior to the germinal center reaction

**DOI:** 10.1101/737908

**Authors:** Francesc Català-Moll, Dieter Weichenhan, Pavlo Lutsik, Ángel F. Álvarez-Prado, Javier Rodríguez-Ubreva, Christian Klemann, Carsten Speckmann, Hassan Abolhassani, Mónica Martínez-Gallo, Romina Dieli-Crimi, Pere Soler-Palacín, Sven Kracker, Lennart Hammarström, Anne Durandy, Bodo Grimbacher, Christoph Plass, Esteban Ballestar

## Abstract

Mutations in activation induced deaminase (AID) lead to hyper-IgM syndrome type 2 (HIGM2), a rare human primary antibody deficiency. AID-mediated cytosine deamination has been proposed as mediating active demethylation, although evidences both support and cast doubt on such a role. We here made use of HIGM2 B cells to investigate direct AID involvement in active DNA demethylation. HIGM2 naïve and memory B cells both display widespread DNA methylation defects, of which approximately 25% of these defects correspond to active events. For genes that undergo active demethylation that is impaired in HIGM2 individuals, we did not observe AID involvement but a participation of TET enzymes. DNA methylation alterations in HIGM2 naïve B cells are related to premature overstimulation of the B-cell receptor prior to the germinal center reaction. Our data supports a role for AID in B cell central tolerance in preventing the expansion of autoreactive cell clones, affecting the correct establishment of DNA methylation patterns.

## INTRODUCTION

Hyper-IgM syndrome type 2 (HIGM2) is a rare primary antibody deficiency characterized by loss-of-function mutations in activation-induced deaminase (AID)^1^, an enzyme required for several crucial steps of B cell terminal differentiation. AID converts deoxycytosines (dCs) into deoxyuracils (dUs), producing dU:dG mismatches that are removed by mismatch repair and base-excision repair^2–6^. Deaminase activity is required for somatic hypermutation (SHM) and class-switch recombination (CSR) of immunoglobulin genes, which are necessary processes for affinity maturation and antibody diversification within the germinal centers^1,7,8^. AID deficiency results in the absence of CSR and SHM, and leads to lymphoid hyperplasia^1^. HIGM2 patients have normal or elevated serum IgM levels with severe reduction of IgG, IgA, and IgE, resulting in considerable susceptibility to bacterial infections^1^.

In addition to its role in CSR and SHM, AID has been proposed as participating in active DNA demethylation through deamination of 5-methylcytosine, leading to a mismatch that is converted to G:C by thymine DNA glycosylase (TDG), followed by base-excision repair. During the past decade, conflicting reports have both supported and discounted such a role for AID (reviewed in^9^). For instance, in three independent studies using B cells from AID-deficient mice, two reported the absence of DNA methylation changes^10,11^, whereas the third study found DNA methylation differences relative to wild type mice^12^. However, methodological aspects could explain the discrepancies between these studies. In parallel, the discovery of alternative enzymatic pathways that lead to *bona fide* active DNA demethylation through ten-eleven translocation methylcytosine dioxygenase (TET)-mediated oxidation of methylcytosines^13–15^ raised more doubts about the possibility that AID redundantly plays such a role. There is currently no consensus about whether AID is involved in mediating DNA demethylation in specific cell contexts.

Whole-genome analysis has shown the occurrence of a vast amount of demethylation associated with B cell differentiation. Changes occur mostly during naïve B cell activation, yielding memory B cells^16,17^ that coincide with the highest peak of AID expression^1,7,18^. Naïve B cells start to proliferate upon activation by antigen encounter. Then they express AID which triggers the secondary diversification of antibodies by SHM and CSR. This is followed by affinity maturation which finally leads to a) a new cycle of SHM or b) terminal differentiation into memory or plasma B cells depending on the affinity of the B cell receptor (BCR) for the cognate antigen^19^.

In this study, we took advantage of the exceptional possibility to address the direct role of AID in active demethylation by comparing the complete DNA methylomes of naïve and memory B cells of HIGM2 patients with those of healthy individuals. By studying two sibling patients with a homozygous mutation for AID that results in a severely truncated enzyme we were able to determine its direct link with DNA methylation defects and infer its catalytic activity in relation to active DNA demethylation.

Our results show that the absence of AID catalytic activity affects DNA methylation in naïve and memory B cells. The majority of the changes observed in the transition from naïve to memory B cells arise from passive demethylation and are linked to late-replicating domains. However, for those potentially associated with active demethylation, we found no evidence of a direct involvement of AID, and our analysis indicates that TET enzymes are responsible for DNA methylation changes in this cell context. The increased DNA demethylation noted in naïve B cells of HIGM2 patients is associated with a premature demethylation of BCR downstream genes prior to the germinal center reaction. Indeed, we found that these changes are related to the expansion of autoreactive clones, which suggests a major role for AID in preventing the expansion of such clones under normal conditions.

## RESULTS

### Study strategy

We obtained peripheral blood from two HIGM2 sibling patients, both with the same homozygous mutation for AID, and two healthy controls. Specifically, the patients carried a deletion (Exon 2 c.22_40del19) that generated a frameshift variant (p.Arg8Asnfs*19) that affects the majority of AID, including its catalytic domain. We inspected the peripheral B cell compartment by flow cytometry. As previously described, HIGM2 patients are characterized by the absence of class-switched memory B cells (CD19^+^CD27^+^IgM^−^IgD^−^, csMBC)^1^. Nevertheless, classic non-class-switched memory B cells (CD19^+^CD27^+^IgM^+^IgD^+^, ncsMBC) and naïve B cells (CD19^+^CD27^−^IgM+IgD+, NBC) are present in patients (Fig. 1a). Under physiological conditions, ncsMBC cells display certain levels of SHM at the immunoglobulin locus^16^, which supports the expression of AID during their maturation in germinal centers^1,7,18^. Therefore, the comparison between the DNA methylation profiles from NBC and ncsMBC of healthy and HIGM2 individuals are an adequate model for testing the potential role of AID in demethylation.

**Figure 1.**
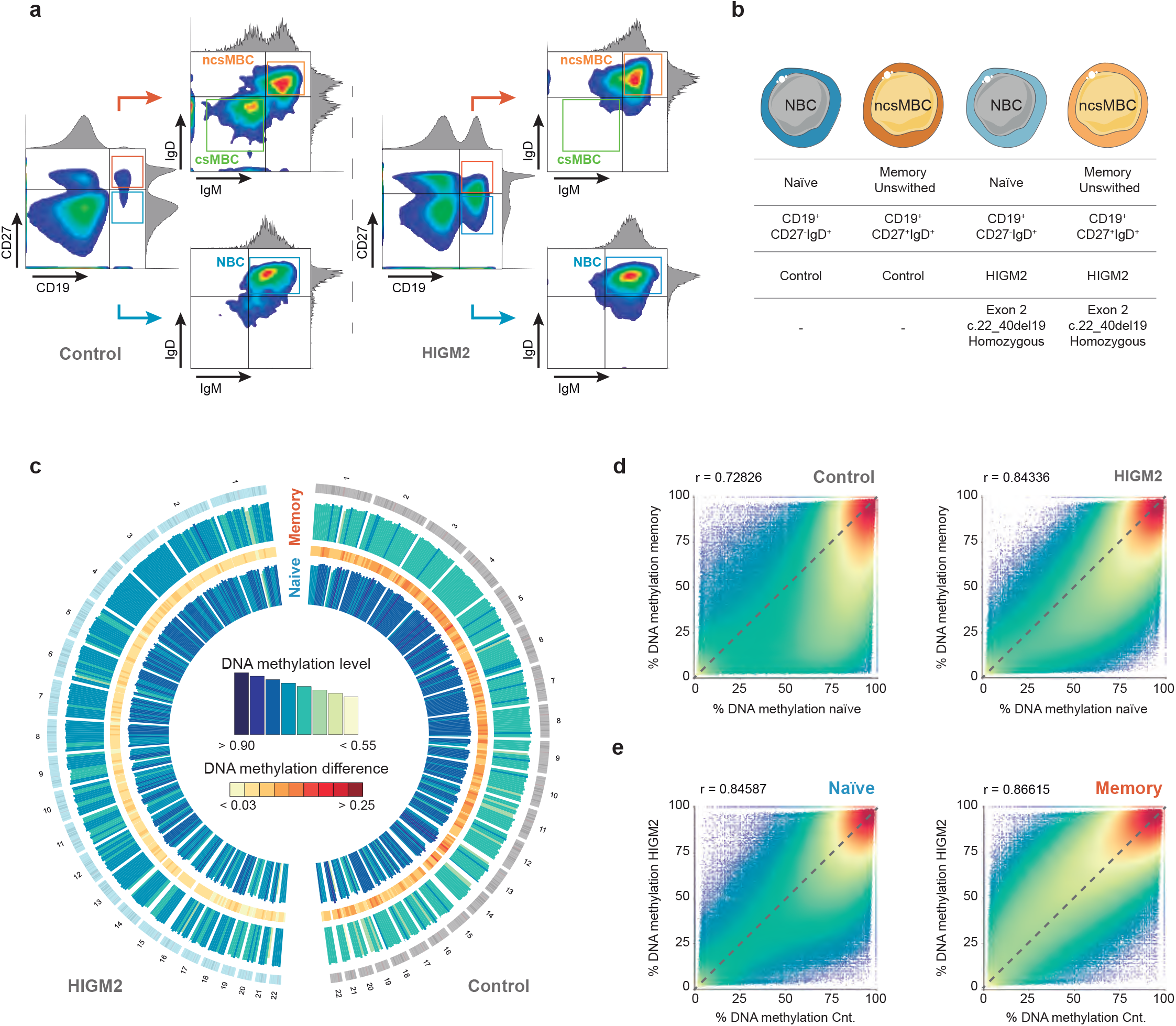
DNA methylome of HIGM2 B cell subpopulations determined by T-WGBS. (**a**) Representative example of strategy for sorting B cell populations (naïve B cells, NBC; classic non-class-switched memory B cells, ncsMBC; class-switched memory B cells, csMBC). (**b**) Description of B cell subpopulations analyzed. (**c**) Circular representation of DNA methylation levels for naïve B cells (inner circle) and unswitched memory B cells (outer circle) for controls (right) and HIGM2 patients (left). Histogram tracks represent the average methylation levels over 10 Mb windows. Heatmap shows the DNA methylation differences between naïve and memory B cells (**d**) Scatter plots with density information and color scale for the pairwise comparisons of control naïve to control memory B cells and HIGM2 naïve to HIGM2 memory B cells. (**e**) Scatter plots with density information and color scale of the pairwise comparisons of HIGM2 naïve to Control naïve B cells and HIGM2 memory to Control memory B cells.

### HIGM2 patients display an aberrant methylation profile in naïve and unswitched memory B cells

We performed tagmentation-based whole-genome bisulfite sequencing (T-WGBS), a version of the WGBS method that allows analysis of limited DNA amounts^20^, for two biological replicates of each of the two aforementioned B cell subsets of the HIGM2 and of healthy controls (from now on referred to as “naïve” and “memory” cells) (Fig. 1b). Pearson correlation and t-distribution stochastic neighbor embedding (t-SNE) between samples was highly reproducible between replicates (correlation coefficient > 0.9, **Supplementary Fig. 1a,b**). We also compared our methylation data from healthy controls with public data from the International Cancer Genome Consortium (ICGC)^21^ and Oakes et al.^16^, thereby confirming the robustness of our data (**Supplementary Fig. 1c,d**).

Global inspection of DNA methylation confirmed that, as reported^16,17^, transition from naïve to memory B cells is accompanied by global demethylation of the genome (Fig. 1c). However, the same comparison in HIGM2 patients showed a partial impairment of demethylation during B cell differentiation (Fig 1c,d), compatible with a potential role of AID as a demethylating enzyme. Unexpectedly, we also observed that naïve B cells were more demethylated in HIGM2 patients than in healthy controls (Fig. 1e). Taken together, these global observations suggest that AID loss not only affects the DNA methylation patterns in the transition from naïve to memory B cells, but also has a significant role in establishing the B cell methylome in earlier stages of development.

### A high proportion of DNA demethylation events identified in HIGM2 are due to passive demethylation of late-replicating domains

Recent studies have shown that a high proportion of the demethylation occurring in cancer and in differentiation processes, are associated with high proliferation rates. Such demethylation takes place in regions known as ‘partially methylated domains’ (PMDs), rather than ‘highly methylated domains (HMDs). PMDs are characterized by late replication, and their demethylation is a passive event, as a result of inefficient DNA remethylation during DNA replication^22–27^. Recent reanalysis of the B cell lineage DNA methylation profiles published by the BLUEPRINT consortium^17^ has shown the occurrence of demethylation of PMDs in the transition towards memory cell and antibody-secreting plasma cells^28^. This highlights how critical it is to separate the analysis of DNA methylation changes produced inside and outside PMDs when examining the occurrence of active demethylation processes to exclude those changes due to passive demethylation.

To address this matter, we examined the overlap of the differentially methylated regions (DMRs) corresponding to all comparisons with the PMDs and HMDs obtained by Zhou et al.^23^. We found that the majority of DMRs overlap with PMDs (72.5%, Fig. 2a). DMRs were therefore classified into two sets: the first comprised those DMRs that coincided with common PMDs (PMD-DMRs), and the second contained all the other DMRs (non-PMD-DMRs). Global inspection of the methylation values showed that the two groups of DMRs had intermediate methylation values in memory B cells, although non-PMD-DMRs had lower methylation levels (Fig. 2b), suggesting the presence of active demethylation events.

**Figure 2.**
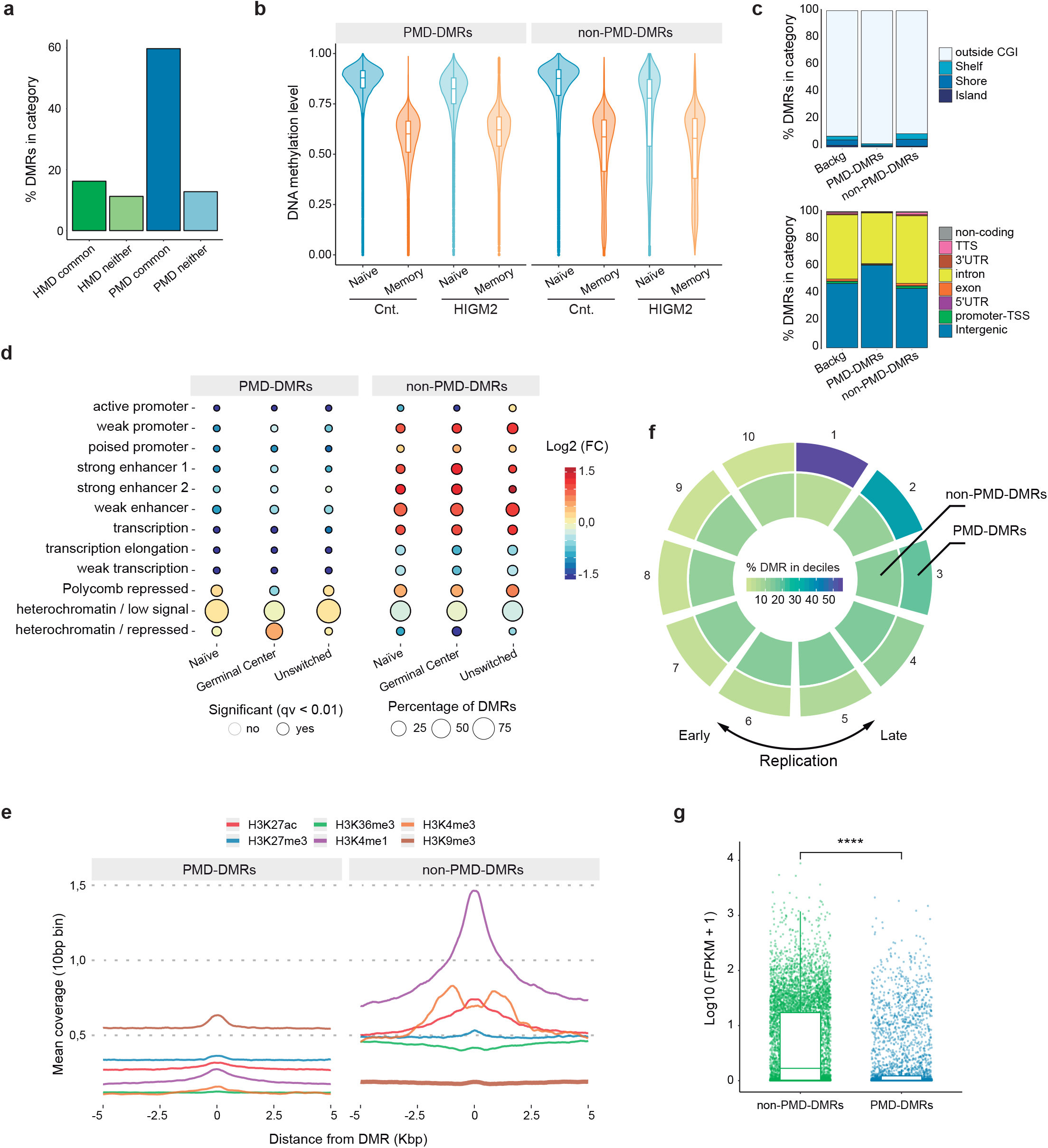
Detection and characterization of partially methylated regions. (**a**) Bar plot showing the percentage of DMRs in PMD-HMD (partially and highly methylated domains, respectively) regions annotated by Zhou and colleagues^23^. ‘Common’ refers to those PMD-HMD regions common to all cell types analyzed by Zhou and colleagues. ‘Neither’ refers to those regions detected only in a fraction of the cell types analyzed. (**b**) Box and violin plots summarizing the distribution of DNA methylation levels per sample group of DMRs inside and outside PMDs. (**c**) Location proportions of PMD-DMRs and non-PMD-DMRs in the context of CpG islands (CGIs) and gene-related regions. (**d**) Bubble chart depicting the enrichment (red) or depletion (blue) of chromatin states. The dot fill represents the logarithmic-fold change, dot size indicates the percentage of DMRs in the chromatin state, and the edge indicates the statistical significance of the enrichment (black: significant, none: not significant; q < 0.01). (**e**) Composite plots of patterns of different histone marks ChIP-seq signals ± 5 kb around the midpoints of PMD-DMRs (left) and non-PMD-DMRs (right) of PMDs. (**f**) Circular representation of the proportion of DMRs in deciles of replication timing data of GM12878 cell line. Color scale represents the proportion of DMRs in each decile. The first and last deciles correspond to the regions of latest and earliest replication, respectively. (**g**) Box and dot plots showing the distribution of expression values of DMRs associated genes inside and outside PMDs. Statistical tests: two-tailed Fisher’s exact (d) and unpaired Wilcoxon’s (g) tests. (FPKM= fragments per kb of transcript per million mapped reads).

Next, we analyzed the functional genomic features of the two groups of DMRs to confirm whether it is appropriate to use the PMD/HMD annotation with our data. We found that the PMD-DMRs were enriched in intergenic regions (Fig. 2c). Considering the connection between the two groups of DMRs and the chromatin states, we found that DMRs occurring in PMDs were mainly associated with heterochromatic regions, while non-PMD-DMRs were highly enriched at enhancers and active promoters, which were mainly associated with active demethylation^29–36^ (Fig. 2d,e).

PMDs are characterized by their association with late-replication domains^22,23^. To test this property in our data, we divided the GM12878 Repli-seq data into deciles and measured the percentage of DMRs in each category. We confirmed that, indeed, the DMRs annotated as PMDs were mainly found in late-replication regions (Fig. 2f) and were accompanied by lower expression levels of associated genes in germinal center B cells (Fig. 2g). Taking all these observations into account, our results indicated that most of the DNA methylation changes in all the comparisons occur in PMDs. However, the existence of a set of non-PMD-DMRs (~27%) located in highly active regions suggests the potential participation of active demethylation events, which could now be interrogated for the potential direct participation of AID.

### HIGM2-associated defects in DNA methylation in the transition from naïve to memory B cells do not have the features of AID targets

Given all the previous considerations, including the removal of DNA methylation changes related to DNA replication (PMD-DMRs), our model allows us to examine whether AID has a direct role in mediating demethylation in the B cell lineage. In this context, germinal center B cells in the transition from naïve to memory, displayed the highest levels of AID mRNA expression (**Supplementary Fig. 2a**)^1,7,37^. This makes the comparison between naïve and memory B cells the most suitable to explore the direct role of AID in active demethylation.

To this end, we selected those DMRs with methylation dynamics consistent with a potential demethylation mediated by AID (P-AID DMRs). These are defined as DMRs that are demethylated in the transition from naïve to memory B cells, without such changes in the HIGM2 patients and having similar methylation levels in naïve cells of controls and HIGM2 patients (Fig. 3a). A total of 522 DMRs (containing 450 different genes) fulfilled these conditions.

**Figure 3.**
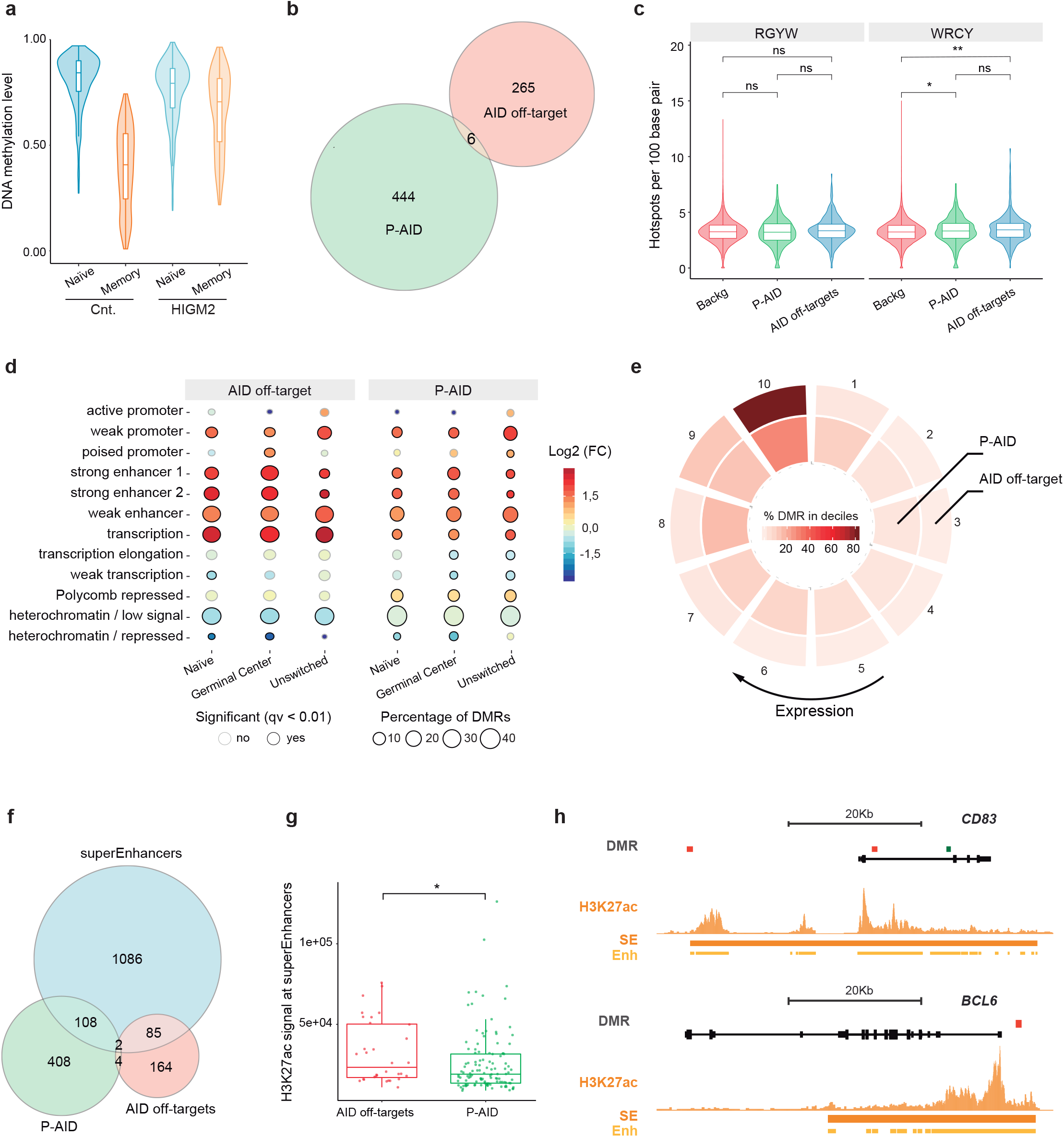
Indirect involvement of AID in DNA demethylation dynamics during B cell activation. (**a**) Box and violin plots summarizing the distribution of DNA methylation levels per sample group of regions potentially demethylated directly by AID (P-AID). (**b**) Venn diagram showing the overlap between P-AID DMRs and human orthologues of the described mouse AID off-target genes. (**c**) Box and violin plots displaying the frequency of WRCY/RGYW (W = A/T; R = G/A; Y = C/T) hot spots per 100 bp in DMRs of each subset. (**d**) Bubble chart depicting the enrichment (red) or depletion (blue) of chromatin states. The dot fill represents the logarithmic-fold change, the dot size shows the percentage of DMRs in the chromatin state, and the edge indicates the statistical significance of the enrichment (black: significant, none: not significant, q < 0.01). (**e**) Circular representation of proportion of DMRs in deciles of germinal center B cells. Color scale represents the proportion of DMRs in each decile. The first and last deciles correspond to the least and most highly expressed genes, respectively. (**f**) Venn diagram of the overlap between P-AID and AID off-target DMRs with germinal center super-enhancers. (**g**) Box plots showing H3K27ac signal of AID off-target (red) and P-AID (green) associated super-enhancers. The one-sided unpaired Wilcoxon’s test was used to examine signal intensity differences. (**h**) DNA methylation and H3K27ac profiles in the vicinity of two representative genes. DMR color indicates the type: P-AID (green) and AID off-target DMRs (red). At the bottom, the H3K27ac ChIP-seq signal in germinal center B cells is shown in dark orange. Identified super-enhancers are depicted below with dark orange bars. Enhancer regions are represented with light orange bars. Statistical tests: two-tailed unpaired Wilcoxon’s (c, g) and Fisher’s exact (d) tests.

We first investigated the overlap between the genes contained in the P-AID DMRs and 271 described off-target AID genes (AID targets outside the locus of immunoglobulins)^38^ (add PMID: 17485517; PMID: 9751748; PMID: 11460166; PMID: 18273020) and found a low correspondence (Fig. 3b). We also tested the presence of described AID hot spots^39^ and found a significant increase for the WRCY hot spot with respect to the background, although the increase seemed too slight to be of biological relevance (Fig. 3c). On the other hand, we found that DMRs associated with off-target AID genes were normally demethylated in HIGM2 patients (**Supplementary Fig. 2b**).

Two recent studies have characterized the genomic and epigenomic features of AID off-target regions. Independently, they found that AID targets regions with convergent transcription from intragenic super-enhancers^40,41^. In this sense, although there were no significant differences regarding gene localization between the DMRs associated with AID off-targets and P-AID DMRs (**Supplementary Fig. 2c**), the former exhibited greater enrichment of enhancer regions (Fig. 3d and Supplementary Fig. 2d) that was associated with more transcriptional activity of associated genes in germinal center B cells (Fig. 3e). We also found that, although the two DMR groups had a similar percentage of overlap with super-enhancers (P-AID 21%, AID off-target 33.3%; Fig. 3f), the super-enhancers of the AID off-targets had a stronger signal for H3K27ac (Fig. 3g,h and Supplementary Fig. 2e) and greater transcriptional activity of their associated genes than P-AID (**Supplementary Fig. S2f**). Finally, we hypothesized that if AID had a role mediating active DNA demethylation, we would expect to see differences in CpG sites in a WRCY context at the super-enhancers of AID off-target genes. However, no such differences were observed (**Supplementary Fig. 2g**). Taken together, our results suggest that, despite the differences in DNA methylation associated with B cell activation between wild type and AID-deficient B cells, such demethylation is not directly associated with AID catalytic activity.

### Demethylation during activation of B cells involves TET family proteins

Our findings ruled out a role for AID in directly mediating DNA demethylation in human B cells and contradicted those of a previous study addressing this hypothesis in mouse B cells. However, the conclusions of that study were based on there having been a significant enrichment with SHM target genes and AID-associated dsDNA breaks (as was also the case in the present study), but with an overlap between the two of less than 10%^12^.

While the removal of methyl groups of cytosines mediated by TET enzymes involves the generation of oxidation intermediates^14,31^, the proposed mechanism of DNA demethylation by AID implies that the 5mC conversion in a thymine could be repaired and replaced with an unmethylated cytosine^9,42^ (Fig. 4a). Given these considerations, we hypothesized that if AID deaminates 5mC and yields thymine in mouse B cells at the population level, AID-dependent demethylation events would be associated with lower levels 5mhC than those that are TET-dependent. To address this possibility, we merged the methylation data of Dominguez and colleagues^12^ with public hydroxy-meDIP-seq data of mouse B cell activation^43^. We selected a set of CpGs demethylated in wt but not in AID^−/−^ mice (mouse potential AID targets, P-AID), a second CpG set that is significantly demethylated under both conditions and more likely to be TET-dependent (positive control), and a third group of CpGs without demethylation (negative control; delta DNA methylation < 0.05) (Fig. 4b). We found that mouse P-AID CpGs overlapped little with the described AID off-target genes (< 8%, Fig. 4c). Next, we determined the hydroxymethylation status of the three groups of CpGs, and found that mouse P-AID CpGs presented similar levels of hydroxymethylation to those that are TET-dependent (positive control), and significantly higher levels than CpGs selected as negative controls (Fig. 4d). Taken together, these results indicate that demethylation changes occurring during B cell maturation are mediated by TET proteins, and if AID has a role it should be only in an indirect manner.

**Figure 4.**
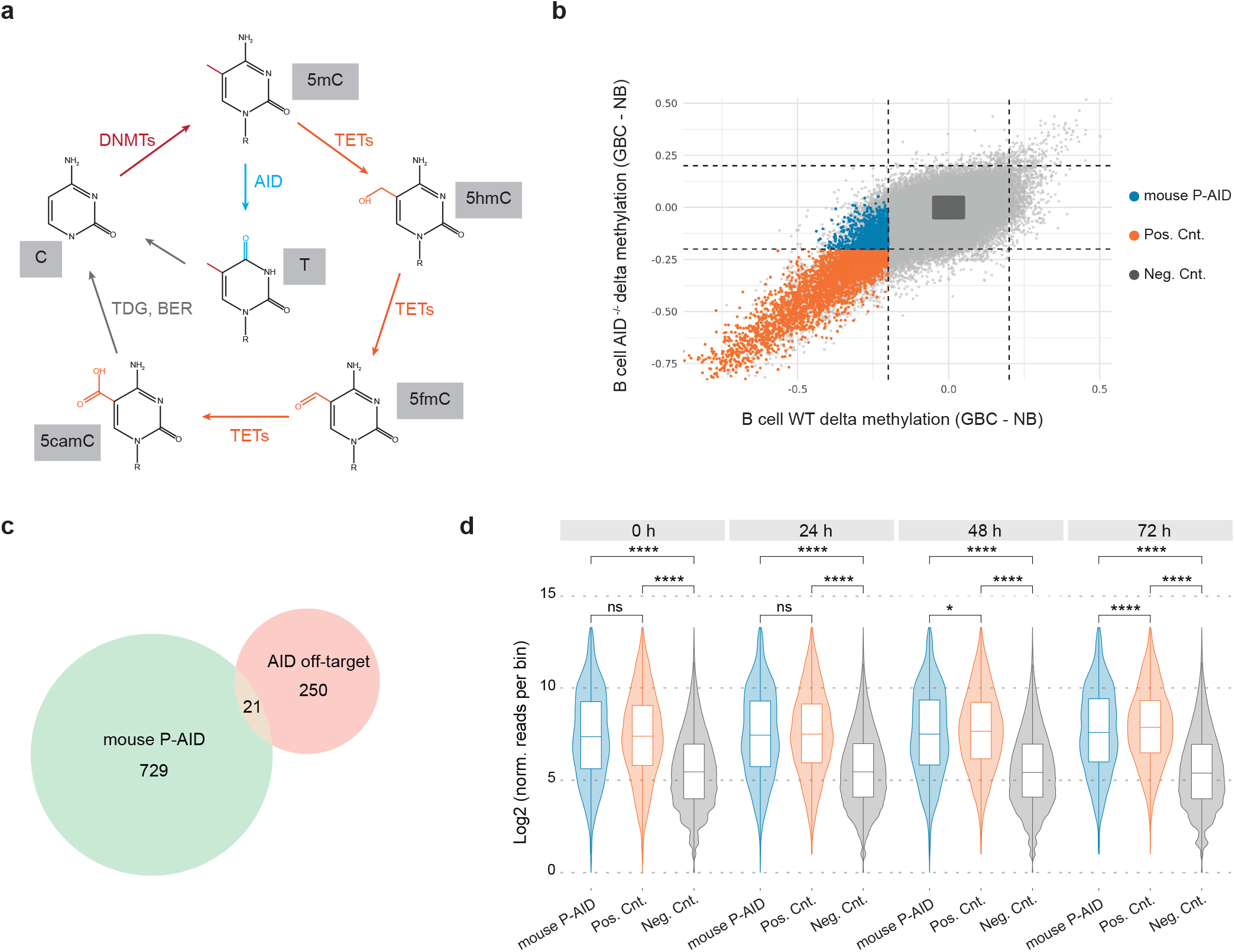
TET proteins are responsible for DNA demethylation events. (**a**) Scheme depicting the potential dynamics of demethylation according to the oxidation recation catañlyzed by TET enzymes or the deamination-mediated reaction catalyzed by AID (**b**) Scatter plot showing a pairwise comparison of DNA methylation differences between naïve and germinal center B cells of WT and AID^−/−^ mouse. Blue dots indicate CpGs potentially demethylated by AID (mouse P-AD). Orange dots indicate those demethylated in both WT and AID DKO (Positive control). Gray dots show those without methylation changes in either condition (Neg. Cnt.). **(c)** Venn diagram showing the overlap between mouse P-AID CpGs and human orthologues of the described mouse AID off-target genes. (**d**) Box and violin plots depicting the distribution of hydroxymethylation values of the sets of CpGs at different times after LPS/IL4 B cell activation. Statistical test: Two-tailed unpaired Wilcoxon’s test (d).

### AID deficiency results in premature demethylation of the BCR pathway of naïve B cells

Our initial analysis suggested that alterations occur in DNA methylation in naïve cells of HIGM2 individuals in comparison with healthy controls. Specifically, HIGM2 naïve B cells appeared to be more demethylated than those of healthy controls. AID expression has customarily been associated with the germinal center reaction^1,7,18^. However, more recent evidence suggests that AID might have a role in earlier stages of B cell development^44–46^.

Comparing HIGM2 and control naïve B cells, we detected 2152 hypomethylated DMRs (Fig. 5a) and 127 hypermethylated DMRs (**Supplementary Fig. 3a**), PMDs having been excluded. Both groups of DMRs were mostly found in intergenic regions and introns (**Supplementary Fig. 3b**). However, while hypomethylation was associated with enhancer regions, hypermethylation was mainly enriched in promoters (**Supplementary Fig. 3c**), in agreement with its reported regulatory role in the ‘spurious’ initiation of transcription^47^.

**Figure 5.**
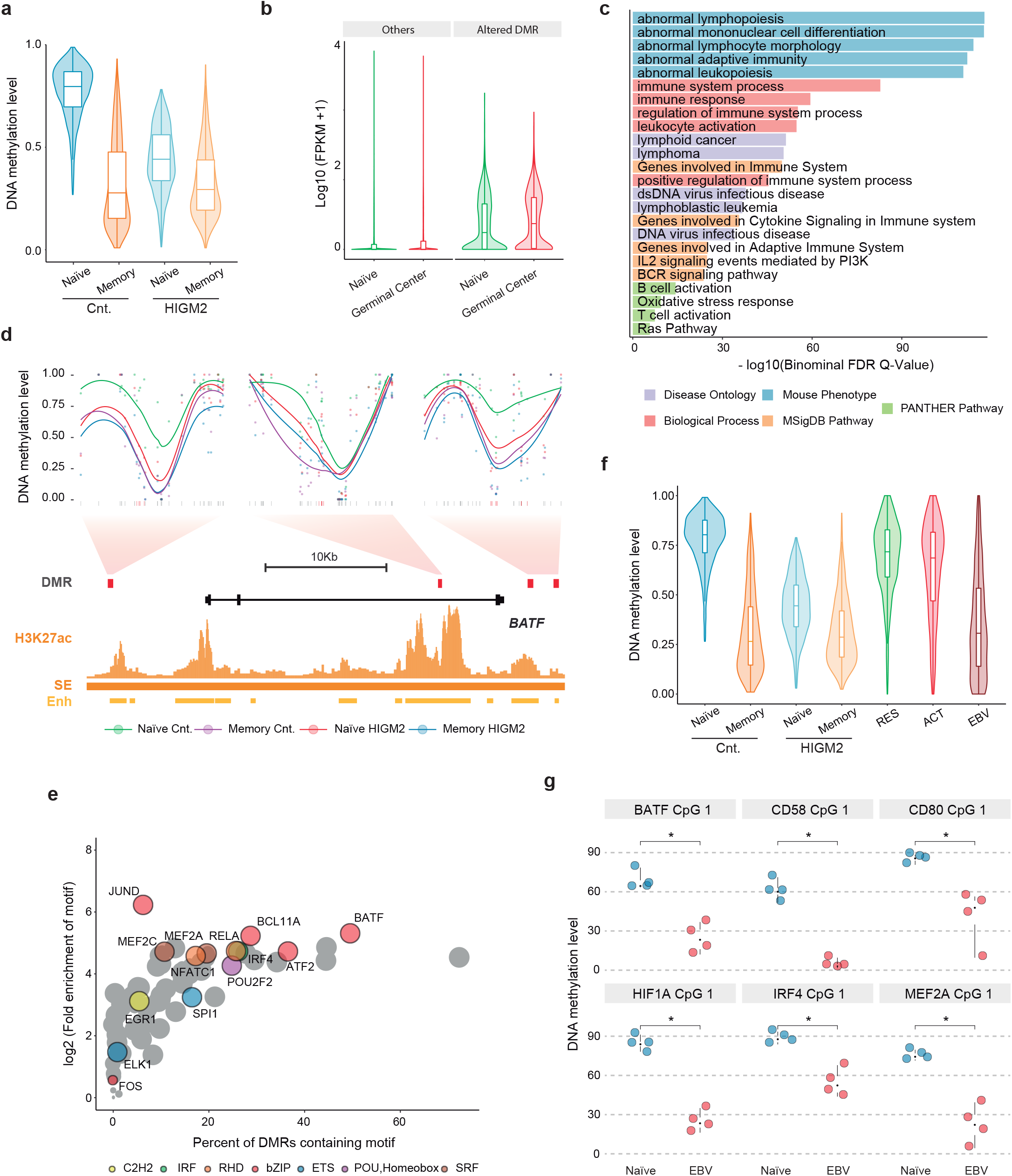
Role of AID in early B cell development. (**a**) Box and violin plots summarizing the distribution of DNA methylation levels per sample group of differentially hypomethylated regions between control and patient naïve B cells. (**b**) Box plot showing the distribution of expression values of altered DMR-associated genes and the all other genes in naïve (green) and germinal center (red) B cells. (**c**) Results of gene set enrichment analysis using GREAT software. The plot depicts the top five enriched terms for five annotation databases, based on *P* values from the binominal distribution. (**d**) Smoothed DNA methylation data of *BATF*-associated altered DMRs and H3K27ac ChIP-seq signal in germinal center B cells. (**e**) Bubble scatter plot of transcription factor ChIP-seq peaks determined in GM12878 lymphoblastoid cell line in altered DMRs. Only transcription factors downstream of BCR signaling are colored according to the transcription factor family. Bubble size corresponds to the logarithm of adjusted values of *P*. (**f**) Box and violin plots summarizing the distribution of DNA hypomethylation levels of altered DMRs in resting B cells (RES), B cells activated with CD40L/IL4 (ACT) and B cells infected with Epstein-Barr virus (EBV). (**g**) Dot plot showing the DNA methylation values determined by pyrosequencing of naïve B cells (naïve B cells) and EBV-infected naïve B cells (EBV). Statistical tests: two-tailed Wilcoxon’s test (g). (FPKM= fragments per kilobase of transcript per million mapped reads, FDR = false discovery rate).

We observed that DMRs in this comparison, i.e. differentially methylated between naïve cells of HIGM2 patients and healthy controls, underwent a similar change in DNA methylation during the transition from naïve to memory cells in controls and HIGM2 patients, suggesting that the DNA methylation alterations in HIGM2 naïve cells correspond to changes that occur later in differentiation, as if these cells were pre-activated outside the germinal center (Fig. 5a). This is consistent with the finding that genes associated with these DMRs became upregulated during the activation of naïve to memory cells in the germinal center (Fig. 5b) and were associated with functional categories related to B cell activation via BCR (Fig. 5c). Some genes that have altered DMRs, such as *BATF* (Fig. 5d)^43,48,49^ or *MEF2A* (**Supplementary Fig. 3d**)^50–52^, are crucial to B cell development.

To explore the possibility that the changes between naïve B cells of HIGM2 patients and controls are due to pre-activation outside the germinal center, we tested the enrichment for transcription factor motifs in the DMRs. Some of the most enriched TFs are downstream of the BCR pathway^53^ (**Supplementary Fig. 3e**). We then validated these results through enrichment analysis of the ChIP-seq data available for GM12878 cells from the ENCODE consortium^54^ and found them to be in accordance with those of the motif enrichment analysis and showed enrichment of downstream TFs of the BCR (Fig. 5e). In addition, some of these TFs were associated with altered DMRs such as BATF, MEF2A, NFATC1, BCL11A and IRF4.

The type III latency state of the Epstein-Barr virus (EBV) is characterized by the constitutive activation of the BCR and CD40 pathways^55–59^, both of which are major signaling pathways that function during B cell activation^60^. In this sense, the B lymphoblastoid cell line GM12878 presents a type III latency state^61^ and is therefore a good model for testing if the changes at the altered DMRs are indeed produced via BCR/CD40. To test this hypothesis, we checked the methylation status of altered DMRs in public DNA methylation data of EBV-transformed B cells and CD40L/IL-4-activated B cells^24^. We observed that EBV transformation effectively reproduced methylation changes of the altered DMRs. Conversely, such changes did not take place with CD40/IL-4 activation, suggesting that the BCR pathway has a significant role (Fig. 5f and Supplementary Fig. 3f). We confirmed these results by transforming naïve B cells with EBV. After 30 days, we used pyrosequencing to test a selection of genes that were aberrantly demethylated in HIGM2 naïve B cells. We found that these genes underwent demethylation following EBV-mediated transformation of naïve B cells (Fig. 5g and Supplementary Fig. 3g). Taken together, our findings showed that the changes between HIGM2 naïve B cells and those of the controls are due to the aberrant pre-activation of the BCR at some point of B cell development prior to the germinal center reaction.

The analysis of motif enrichment and ChIP-seq data in GM12878 indicated that BATF is one of the main potential mediator of demethylation by TET proteins via their recruitment in the generation of altered DMRs in HIGM2 naïve B cells (~50% overlap with the DMRs). BATF is a regulator of B and T cell activation (BCR and TCR pathways, respectively), in cooperation with IRF4^48,62,63^. We observed that 87% of the ChIP-seq peaks of IRF4 overlapped with BATF peaks in GM12878 (Fig. 6a). As expected, we were able to confirm that regions with BATF and IRF4 binding had lower DNA methylation levels in naïve cells of HIGM2 patients than in controls (Fig. 6b), as well as JUND, another TF downstream of BCR (**Supplementary Fig 4a**). This did not occur in regions enriched for other B cell-intrinsic TF binding motifs (**Supplementary Fig. 4a**). Using mRNA transcription data from IRF4 and BATF knockouts from GM12878 cells^61^, we determined that genes with binding motifs for both TFs displayed expression changes for many of these genes associated with altered DMRs (Fig. 6c). All these results suggest that a significant fraction of the altered DMRs in HIGM2 naïve B cells may be associated with the recruitment of the BATF/IRF4 complex to these genomic sites. Such recruitment might facilitate TET protein-mediated hydroxymethylation and the consequent demethylation of those regions, as recently reported^43^.

**Figure 6.**
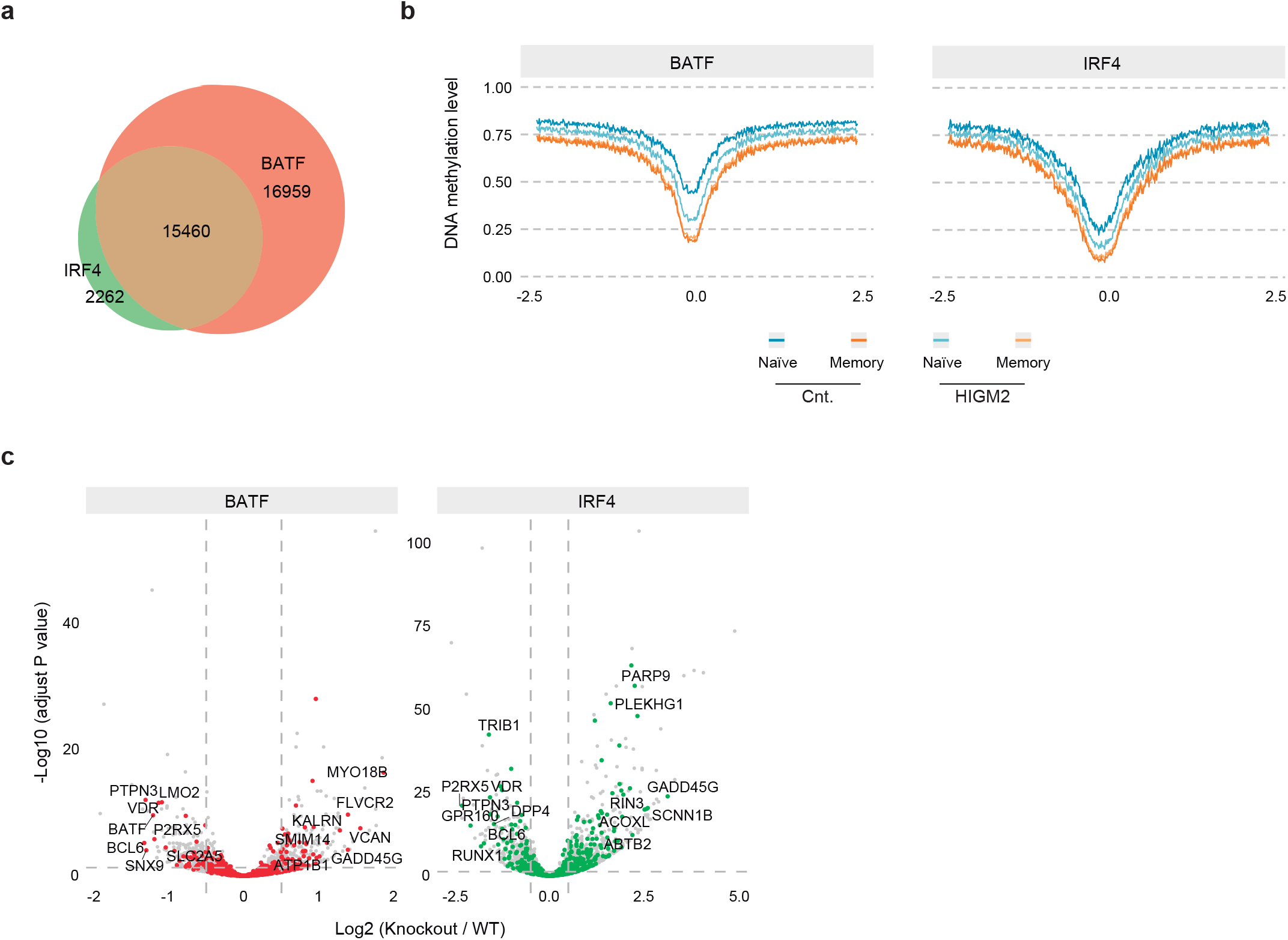
BATF / IRF4 complex-mediated part of methylation events. (**a**) Venn diagram representing the overlap between BATF and IRF4 ChIP-seq peaks in the GM12878 cell line. (**b**) Composite CpG methylation levels surrounding transcription factor ChIP-seq peaks (2.5 kb). (**d**) Volcano plot showing the effect of the knockout of IRF4 or BATF at the transcriptional level of genes with co-binding of IRF4 and BATF. Dot colors indicates whether the gene has an altered DMR.

### AID deficiency causes blockade of central B cell tolerance with an expansion of pre-activated autoreactive B cells

Our results suggest that naïve B cells are pre-activated in AID deficient patients. However, the stage of B cell differentiation at which this alteration is produced remains to be established. Two independent studies reported a potential role for AID in removing autoreactive B cells during the central B cell tolerance process in bone marrow. Specifically, the immature B cells with auto-reactive BCR were activated and went into a secondary receptor editing process with an increase in AID and recombination-activating gene 2 (RAG2). However, if the autoreactive BCR did not lose self-antigen affinity the genomic instability induced by the overexposure to high levels of AID led to apoptosis. However, AID deficiency reduces the genomic damage that causes expansion of autoreactive B cells^44,45^. With that in mind, we hypothesized that the presence of naïve B cells with a pre-activation methylation signature in HIGM2 patients is a consequence of the impairment of central B cell tolerance that causes autoreactive naïve B cells to accumulate. In fact, it has been reported that 21% of HIGM2 patients suffer some kind of autoimmune disease^64^.

To assess this hypothesis, we first checked whether there is an expansion of naïve autoreactive B cells in HIGM2 patients with respect to controls. To this end, we used a commercial antibody against 9G4^+^ IgG used to detect autoreactive clones in autoimmune diseases like systemic lupus erythematosus and rheumatoid arthritis^65–68^. We observed an expansion the naïve B cell compartment in HIGM2 patients in comparison with healthy controls (Fig. 7a,b). We did not found an expansion of 9G4^+^ in HIGM2 patients with respect to controls (Fig. 7c). However, we observed a significant increase of mean fluorescence intensity for 9g4 staining (Fig. 7a,d), as well as, an expansion of high 9g4^+^ naïve B cells (Fig. 7e). Next, we determined the methylation status of a selection of genes by pyrosequencing of 9G4^−^ (non-autoreactive) and 9G4^+^ (autoreactive) naïve B cells, and found that autoreactive B cells had lower levels of DNA methylation than their non-autoreactive counterparts (Fig. 7f). Overall, our results suggest that the demethylation in naïve B cells of HIGM2 patients compared with control donors is associated with an expansion of pre-activated autoreactive naïve B cells as a consequence of central B cell tolerance impairment mediated by AID deficiency.

**Figure 7.**
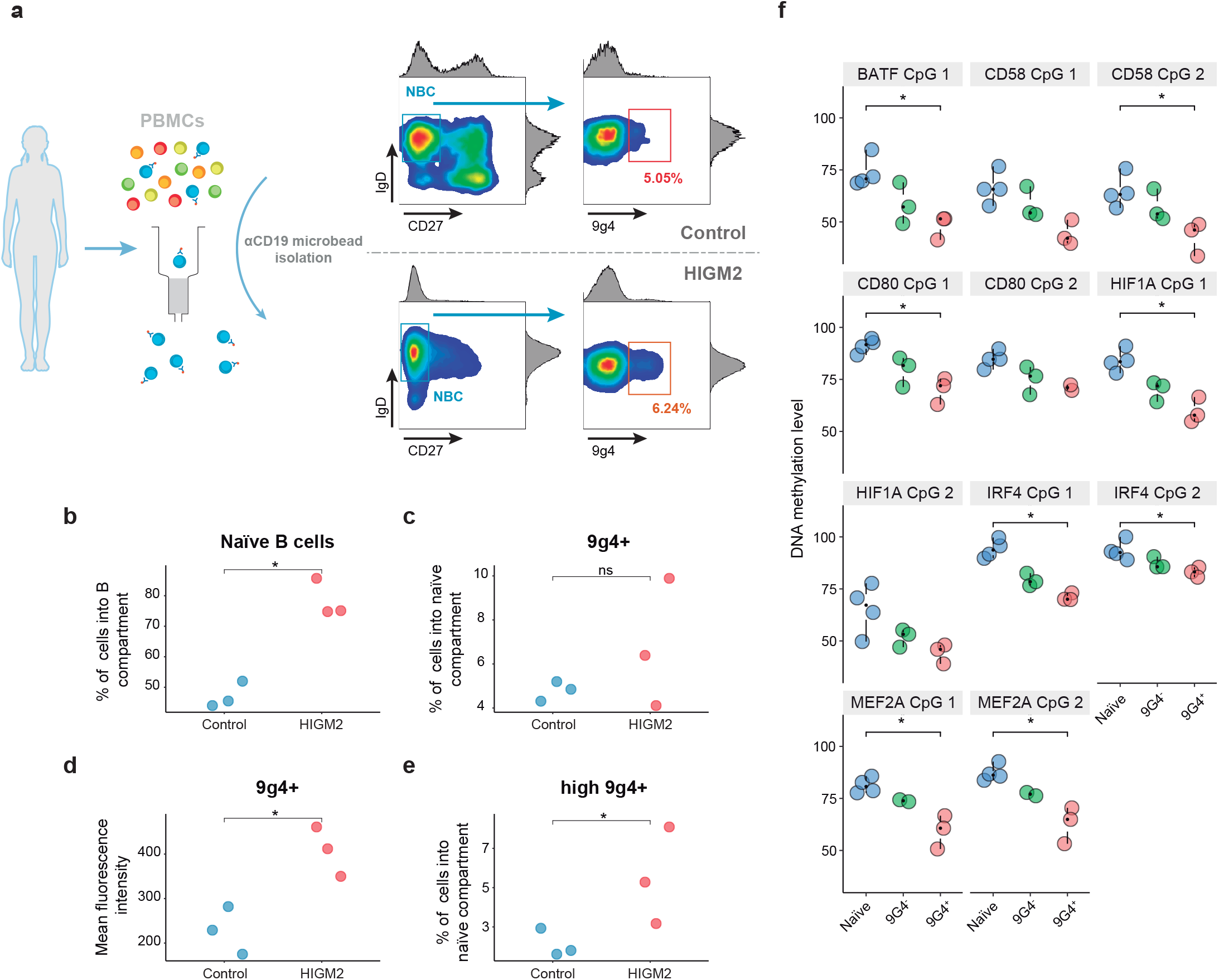
Central B cell tolerance impairment by AID deficiency. (**a**) Representative example of strategy for sorting naïve autoreactive B cells. (**b**) Dot plot showing the percentage of naïve B cells into B cell compartment in HIGM2 patients and controls. (**c**) Dot plot showing the percentage of autoreactive naïve B cells into naïve B cell compartment in HIGM2 patients and controls. (**d**) Dot plot showing the mean fluorescence of 9g4^+^ gate. (**e**) Dot plot showing the percentage of high 9g4^+^ naïve B cells into naïve B cell compartment in HIGM2 patients and controls. (**f**) Dot plot showing the DNA methylation values determined by pyrosequencing of control naïve B cells (naïve), HIGM2 patient 9g4^−^ naïve B cells (9g4^−^) and HIGM2 patient 9g4^+^ naïve B cells (9g4^+^).

## DISCUSSION

Our results show that AID deficiency in HIGM2 syndrome results in the acquisition of aberrant DNA methylation profiles in naïve and memory B cells. Two major conclusions emerge from the study of this phenomenon. First, the analysis of the HIGM2-associated alterations occurring in the transition from naïve to memory B cell rules out the direct involvement of AID in active demethylation. Second, the comparison of naïve B cells in HIGM2 and healthy controls shows premature demethylation of genes downstream of the BCR in AID-deficient individuals, whichis associated with the expansion of autoreactive B cell clones, prior to the germinal center reaction. This reinforces a novel role for AID in preventing the expansion of autoreactive B cell clones, affecting the DNA methylation profiles of naïve B cells.

Our study unequivocally demonstrates that AID does not play a direct role through its catalytic activity in mediating active demethylation in the transition from naïve to memory B cells. This transition is that associated with the highest proportion of DNA methylation changes of the entire B cell differentiation process^16,17^ and also coincides with the highest peak of AID expression. Previous studies addressing the potential participation of AID in demethylation had considered late-replicating domains or the relationship between DNA methylation changes and the genomic features of AID targets. In our study, we determined that most of the changes taking place during the transition from naïve to memory B cells occur through passive demethylation. No associations with AID targets were found for the other changes. These findings are also in line with those of Álvarez-Prado and colleagues^38^, who have indicated that AID-mediated mutation frequencies are too low, In this sense, such low frequency but we unlikely to produce a perceptible effect at the level of DNA methylation.

A second major conclusion of our study concerns the identification of DNA methylation defects in naïve B cells from HIGM2 patients in relation to healthy controls. Customarily, AID expression has been regarded as being restricted to germinal center B cells, but some evidence suggests that AID may also have a role in central B cell tolerance^44,45,69^. In keeping with this, during B cell development, these cells not only become activated in the germinal center but also in previous stages of differentiation in the bone marrow. In that location, self-reactive immature B cells are activated in a process characterized by the upregulation of both AID and recombination-activating gene 2 (RAG2) and the downregulation of the anti-apoptotic MCL-1^70–72^. In this context, AID activity increases the probability of genomic damage with the subsequent activation of apoptosis through p53, which is also enhanced by the inhibition of the anti-apoptotic proteins BCL2 and MCL-1^73^. At that point, self-reactive immature B cells that are unable to correct their affinity for self-antigens by receptor editing are eliminated. In patients with AID deficiency, this mechanism of cell removal is impaired and autoreactive cells accumulate^44^. Indeed, we noted an accumulation of autoreactive B cells in HIGM2 patients than in healthy donors, a finding that is compatible with the previously described high frequency of autoimmune disorders in this type of patient^74^. The failure in AID function in these patients could be responsible for the smaller degree of genomic damage that promotes the expansion of autoreactive naïve B cells. These self-reactive B cells, owing to the persistent activation of their BCR during negative selection in the bone marrow, display a more demethylated profile in genes downstream of the BCR compared with non-autoreactive naïve B cells. Our results therefore indicate that the enhanced demethylation of BCR downstream targets in HIGM2 naïve B cells may be the result of the expansion of autoreactive B cell clones as a consequence of the absence of AID. Our results are also consistent with the recent observation that class-switch recombination occurs infrequently in germinal centers ^75^.

## METHODS

### Human samples

Patients who fulfilled the diagnostic criteria for hyper-IgM syndrome type 2 were included in the study based on ESID clinical diagnostic criteria^76^ and genetic confirmation of *AICDA* mutation and exclusion of other primary and secondary causes of immunodeficiencies. Samples come from the Medical Center of the University Hospital, University of Freiburg, Freiburg, Germany and Hospital Universitari Vall d’Hebron, Barcelona, Spain. The Committees for Human Subjects of the local hospitals approved the study, which was conducted in accordance with the ethical guidelines of the 1975 Declaration of Helsinki. All samples were in compliance with the guidelines approved by the local ethics committee and all donors (and/or their parents) received oral and written information about the possibility that their blood would be used for research purposes.

### Isolation of B cell populations

Peripheral blood mononuclear cells (PBMCs) were obtained from blood. After Ficoll-Isopaque density centrifugation (Rafer, Zaragoza, Spain), collected cells were washed twice with ice-cold PBS, followed by centrifugation at 2000 rpm for 5 min. Next, cells were labeled with antibodies to CD19 – FITC (Miltenyi Biotec, clone LT19), CD27 – APC (Miltenyi Biotec, clone M-T271), IgD – PE (SouthernBiotech, Cat. No. 2032-09) and IgM – PerCP/Cy5.5 (BioLegend, clone MHM-88) for 20 min on ice in staining buffer (PBS with 4% FBS and 2 mM EDTA). Naïve B cells (CD19^+^ CD27^−^ IgD^+^) and unswitched memory B cells (CD19^+^ CD27^+^ IgD^+^) were obtained by FACS sorting on a MoFlo Astrios (Beckman Coulter). Purified samples were pelleted and stored at −80°C.

For isolation of naïve autoreactive B cells. Total B cells were isolated from PBMCs using positive selection with MACS CD19 microbeads (Miltenyi Biotec). Next, cells were stained with CD27-APC (Miltenyi Biotec, clone M-T271), IgD – PE (SouthernBiotech, Cat. No. 2032-09), HLA-DR – PE-Cy7 (eBioscience, clone LN3), 9g4 primary ab (igm Bioscience) and donkey ant-rat IgG (H + L) – Alexa Fluor 488 (invitrogen). 9g4+ naïve B cells (CD27^−^ IgD^+^ 9g4^+^) and 9g4-naïve B cells (CD27^−^ IgD^+^ 9g4^−^) were obtained by FACS sorting on a BD FACSAria II (BD Biosciences). Purified samples were pelleted and stored at −80°C.

### Genomic DNA extraction

For whole-genome bisulfite sequencing, DNA was extracted with a QIAamp DNA micro kit (Qiagen) according to the manufacturer’s protocol. For pyrosequencing experiments, DNA was extracted with a Maxwell RSC Cultured Cells DNA kit (Promega).

### Tagmentation-based whole-genome bisulfite sequencing

For whole-genome bisulfite sequencing, 30 ng of genomic DNA was used to produce four independent barcoded sequencing libraries per DNA sample using the tagmentation method^20^. Sequencing of the TWGBS libraries was done on a HiSeq 2000, PE 125 bp mode. Bisulfite sequencing reads were processed by the DKFZ bisulfite analysis workflow. In brief, the reads were trimmed using Trimmomatic, pre-processed and aligned using MethylCTools, with default parameters (V. Hovestadt, S. Picelli, B. Radlwimmer, M.Z. and P.L., unpublished data), which uses the Burrows-Wheeler alignment algorithm^77^. Following quality control of bisulfite conversion (>99.5% in all samples) and of read-mapping (80-90% could be mapped on average), we performed methylation calling using methylCtools. A summary of the sequencing data for each sample is provided in Supplementary Table 1

### DMR calling

Differentially methylated regions were detected with the DeNovoDMR algorithm included in the Specific Methylation Analysis and Report Tool (SMART2)^78^ using all the default parameters except for the segment CpG number threshold, which was set to 4, the absolute mean methylation difference, which was set to 0.2, and a threshold value of *P* of 0.01. Only those CpGs with a coverage of ≥5 in all samples were considered in the construct of the SMART input matrix. DMR calling was performed or all possible comparisons between naïve and memory B cells for both control and HIGM2 patients.

### Bisulfite pyrosequencing

500 ng of genomic DNA was converted with an EZ DNA Methylation-Gold kit (Zymo Research), following the manufacturer’s instructions. Bisulfite-treated DNA was PCR amplified using primers (see Supplementary Table 2) designed with PyroMark Assay Design 2.0 software (Qiagen). Finally, PCR amplicons were pyrosequenced with the PyroMark Q24 system and analyzed with PyroMark CpG software (Qiagen).

### ChIP-seq data processing

Sequencing reads from ChIP-seq experiments from the BLUEPRINT consortium^79^ were mapped to the hg19 assembly of human reference genome using Burrows-Wheeler Aligner (BWA) v0.7.13^77^ (with parameters -q 5, -l 32, -k 2). After removing reads with MAPQ < 30 with Sequence Alignment/Map (SAMtools) v1.2^80^, PCR duplicates were eliminated using the Picard function available in MarkDuplicates software v1.126^81^. Peak calling was performed using macs2^82^ (with parameters -p 1e-2 --nomodel --shift 0 -B --SPMR). Only peaks with an overlap of ≥ 0.5 between replicates were considered. Histone mark signals around DMR sets were extracted with the annotatePeaks.pl algorithm available in Hypergeometric Optimization of Motif EnRichment (HOMER) software v4.10.3 (with parameters: size = 10000, hist = 10).

### Super-enhancer identification

H3K27ac ChIP-seq data were used to identify the super-enhancer regions, as described previously^83^ using Rank-Ordering of Super-Enhancers (ROSE) software. An enhancer stitching distance of 15 kb was used along with a 2.5 kb transcriptional start site (TSS)-exclusion window.

### Data analysis

Hierarchical clustering was carried out based on Pearson correlation distance metrics and average linkage criteria. For low-dimensional analysis we used the t-distributed stochastic neighbor embedding (t-SNE) method implemented in the Rtsne v0.15 package.

Transcription factor motifs were enriched for each set of DMRs using HOMER software v4.10.3. Specifically, we used the findMotifsGenome.pl algorithm (with parameters -size given -cpg) to search for significant enrichment against a background sequence adjusted to have similar CpG and GC contents.

Transcription factor binding analysis was performed interrogating the overlap between the different sets of DMRs with ChIP-seq data for transcription factors available for GM12878 cell line from the ENCODE Project^54^. The enrichment factor was calculated against random regions as a background, and *P* values were calculated using Fisher’s exact test. Finally, the transcription factors downstream of the BCR signaling pathway were manually annotated from a curated database^53^.

Chromatin states and histone mark enrichments analysis for NBC, germinal center B cells and ncsMBC were assessed using a custom adaptation of the EpiAnnotator R package^84^ using BLUEPRINT data^79^. DMRs were converted to hg38 assembly with the liftOver function in the rtracklayer v1.42 R package.

Replication timing data in the GM12878 lymphoblastoid cell line were obtained from the UW Repli-seq track of the UCSC Genome Browser. Replication timing values were binned in deciles to perform the overlap with the DMR groups.

DMR annotation for genetic context location was performed using the annotatePeaks.pl algorithm in the HOMER software v4.10.3. For determine the location relative to a CpG island (CGI), we used ‘hg19_cogs’ annotation in the annotatr v1.8 R package.

GREAT software^85^ was used to enrich downstream pathways and gene ontologies. We used the single nearest gene option for the association between genomic regions with genes.

All statistical analysis (excluding T-WGBS and ChIP-seq analyses) were done in R v3.5.1. Data distributions were tested for normality. Normal data were tested using two-tailed unpaired Student’s t-tests; non-normal data were analyzed with the appropriate non-parametric statistical test. Levels of significance are indicated as: *, P < 0.05; **, P < 0.01; ***, P < 0.001; ****, P < 0.0001. Non-significance (P ≥ 0.05) was indicated as ‘ns’.

### Public RRBS of B cell activation

Data of EBV and CD40/IL4 B cell activation were downloaded from the NCBI Gene Expression Omnibus (GSE49629)^24^. Methylation calls from RRBS data were filtered, so that only those CpGs with a minimum of five reads per position in all samples were retained. Since RRBS genomic coverage is significantly lower than T-WGBS we only tested the methylation status of positions common to two datasets.

### EBV infection

For naïve B cell EBV infection experiments, we obtained buffy coats from anonymous donors through the Catalan Blood and Tissue Bank (CBTB). The CBTB follows the principles of the World Medical Association (WMA) Declaration of Helsinki. Before providing the first blood sample, all donors received detailed oral and written information and signed a consent form at the CBTB. PBMCs were isolated using Ficoll-Paque gradient centrifugation. Total B cells were isolated from PBMCs using positive selection with MACS CD19 microbeads (Miltenyi Biotec). Next, cells were stained with CD27-APC (Miltenyi Biotec, clone M-T271) and IgD – PE (SouthernBiotech, Cat. No. 2032-09) and naïve B cells were sorted as CD27^−^IgD^+^. Pure naïve B cells were incubated with B95-8 cell supernatant for 3 h at 37ºC in order to infect them with EBV. Finally, cells were collected after 30 days.

## DATA AVAILABILITY

The data that support the findings of this study are available from the NCBI Gene Expression Omnibus (GEO).

## CODE AVAILABILITY

Code and data processing scripts are available from the corresponding author upon request

## FUNDING

We thank CERCA Programme/Generalitat de Catalunya for institutional support. E.B. was funded by the Spanish Ministry of Economy and Competitiveness (MINECO; grant numbers SAF2014-55942-R and SAF2017-88086-R) and was cofunded by FEDER funds/ European Regional Development Fund (ERDF) – a way to build Europe. E.B is supported by RETICS network grant from ISCIII (RIER, RD16/0012/0013).

## ACKNOWLEDGEMENTS

This study makes use of data generated by the BLUEPRINT Consortium. A full list of the investigators who contributed to the generation of the data is available from www.blueprint-epigenome.eu. Funding for the project was provided by the European Union’s Seventh Framework Programme (FP7/2007–2013) under grant agreement no 282510—BLUEPRINT. We are also very grateful to Dr Javier di Noia, Dr Almudena Ramiro, Dr Javier Carmona and Dr Biola M. Javierre for useful feedback.

## CONTRIBUTIONS

F.C.-M. designed and performed experimental experiments and bioinformatics analysis; F.C.-M., D.W. and C.P. generated TWGBS data; P.L. supervised some bioinformatics analysis; F.C.-M and A.F.Á.-P performed AID hot spot analysis; F.C.-M and J.R.-U. design sorting strategies; C.K., C.S, H.A., M.M.-G, R.D., P.S.-P., S.K., L.H., A.D., and B.D. contributed with clinical material and clinical interpretation of the results; E.B. conceived and supervised the study; F.C.-M. and E.B. wrote the manuscript; All authors participated in discussions and interpretation of the data and results.

## REFERENCES

1. Revy, P. et al. Activation-induced cytidine deaminase (AID) deficiency causes the autosomal recessive form of the Hyper-IgM syndrome (HIGM2). Cell 102, 565–75 (2000).

2. Poltoratsky, V., Goodman, M. F. & Scharff, M. D. Error-prone candidates vie for somatic mutation. J. Exp. Med. 192, F27–30 (2000).

3. Di Noia, J. & Neuberger, M. S. Altering the pathway of immunoglobulin hypermutation by inhibiting uracil-DNA glycosylase. Nature (2002). doi:10.1038/nature00981

4. Rada, C., Ehrenstein, M. R., Neuberger, M. S. & Milstein, C. Hot spot focusing of somatic hypermutation in MSH2-deficient mice suggests two stages of mutational targeting. Immunity (1998). doi:10.1016/S1074-7613(00)80595-6

5. Rada, C. et al. Immunoglobulin isotype switching is inhibited and somatic hypermutation perturbed in UNG-deficient mice. Curr. Biol. (2002). doi:10.1016/S0960-9822(02)01215-0

6. Rada, C., Di Noia, J. M. & Neuberger, M. S. Mismatch recognition and uracil excision provide complementary paths to both Ig switching and the A/T-focused phase of somatic mutation. Mol. Cell (2004). doi:10.1016/j.molcel.2004.10.011

7. Muramatsu, M. et al. Class switch recombination and hypermutation require activation-induced cytidine deaminase (AID), a potential RNA editing enzyme. Cell (2000). doi:10.1016/S0092-8674(00)00078-7

8. Arakawa, H., HauschiLd, J. & Buerstedde, J. M. Requirement of the activation-induced deaminase (AID) gene for immunoglobulin gene conversion. Science (80-.). (2002). doi:10.1126/science.1067308

9. Ramiro, A. R. & Barreto, V. M. Activation-induced cytidine deaminase and active cytidine demethylation. Trends in Biochemical Sciences (2015). doi:10.1016/j.tibs.2015.01.006

10. Fritz, E. L. et al. A comprehensive analysis of the effects of the deaminase AID on the transcriptome and methylome of activated B cells. Nat. Immunol. (2013). doi:10.1038/ni.2616

11. Hogenbirk, M. A. et al. Differential Programming of B Cells in AID Deficient Mice. PLoS One (2013). doi:10.1371/journal.pone.0069815

12. Dominguez, P. M. et al. DNA Methylation Dynamics of Germinal Center B Cells Are Mediated by AID. Cell Rep. (2015). doi:10.1016/j.celrep.2015.08.036

13. Kriaucionis, S. & Heintz, N. The nuclear DNA base 5-hydroxymethylcytosine is present in purkinje neurons and the brain. Science (80-.). (2009). doi:10.1126/science.1169786

14. Tahiliani, M. et al. Conversion of 5-methylcytosine to 5-hydroxymethylcytosine in mammalian DNA by MLL partner TET1. Science (80-.). (2009). doi:10.1126/science.1170116

15. Ito, S. et al. Role of tet proteins in 5mC to 5hmC conversion, ES-cell self-renewal and inner cell mass specification. Nature (2010). doi:10.1038/nature09303

16. Oakes, C. C. et al. DNA methylation dynamics during B cell maturation underlie a continuum of disease phenotypes in chronic lymphocytic leukemia. Nat. Genet. (2016). doi:10.1038/ng.3488

17. Kulis, M. et al. Whole-genome fingerprint of the DNA methylome during human B cell differentiation. Nat. Genet. 47, 746–756 (2015).

18. Muramatsu, M. et al. Specific expression of activation-induced cytidine deaminase (AID), a novel member of the RNA-editing deaminase family in germinal center B cells. J. Biol. Chem. (1999). doi:10.1074/jbc.274.26.18470

19. Victora, G. D. & Nussenzweig, M. C. Germinal Centers. Annu. Rev. Immunol. (2012). doi:10.1146/annurev-immunol-020711-075032

20. Wang, Q. et al. Tagmentation-based whole-genome bisulfite sequencing. Nat. Protoc. (2013). doi:10.1038/nprot.2013.118

21. Hudson, T. J. et al. International network of cancer genome projects. Nature (2010). doi:10.1038/nature08987

22. Berman, B. P. et al. Regions of focal DNA hypermethylation and long-range hypomethylation in colorectal cancer coincide with nuclear laminag-associated domains. Nat. Genet. (2012). doi:10.1038/ng.969

23. Zhou, W. et al. DNA methylation loss in late-replicating domains is linked to mitotic cell division. Nat. Genet. (2018). doi:10.1038/s41588-018-0073-4

24. Hansen, K. D. et al. Large-scale hypomethylated blocks associated with Epstein-Barr virus-induced B-cell immortalization. Genome Res. (2014). doi:10.1101/gr.157743.113

25. Lister, R. et al. Human DNA methylomes at base resolution show widespread epigenomic differences. Nature 462, 315–22 (2009).

26. Timp, W. et al. Large hypomethylated blocks as a universal defining epigenetic alteration in human solid tumors. Genome Med. (2014). doi:10.1186/s13073-014-0061-y

27. Robinson, W. P. & Price, E. M. The human placental methylome. Cold Spring Harb. Perspect. Med. (2015). doi:10.1101/cshperspect.a023044

28. Durek, P. et al. Epigenomic Profiling of Human CD4+ T Cells Supports a Linear Differentiation Model and Highlights Molecular Regulators of Memory Development. Immunity (2016). doi:10.1016/j.immuni.2016.10.022

29. Bogdanović, O. et al. Active DNA demethylation at enhancers during the vertebrate phylotypic period. Nat. Genet. (2016). doi:10.1038/ng.3522

30. Neri, F. et al. Single-Base resolution analysis of 5-formyl and 5-carboxyl cytosine reveals promoter DNA Methylation Dynamics. Cell Rep. (2015). doi:10.1016/j.celrep.2015.01.008

31. Lu, F., Liu, Y., Jiang, L., Yamaguchi, S. & Zhang, Y. Role of Tet proteins in enhancer activity and telomere elongation. Genes Dev. (2014). doi:10.1101/gad.248005.114

32. Hon, G. C. et al. 5mC oxidation by Tet2 modulates enhancer activity and timing of transcriptome reprogramming during differentiation. Mol. Cell (2014). doi:10.1016/j.molcel.2014.08.026

33. Rasmussen, K. D. & Helin, K. Role of TET enzymes in DNA methylation, development, and cancer. Genes and Development (2016). doi:10.1101/gad.276568.115

34. An, J. et al. Acute loss of TET function results in aggressive myeloid cancer in mice. Nat. Commun. (2015). doi:10.1038/ncomms10071

35. Yamazaki, J. et al. TET2 mutations affect Non-CpG island DNA methylation at enhancers and transcription factor-binding sites in chronic myelomonocytic Leukemia. Cancer Res. (2015). doi:10.1158/0008-5472.CAN-14-0739

36. Rasmussen, K. D. et al. TET2 binding to enhancers facilitates transcription factor recruitment in hematopoietic cells. Genome Res. gr.239277.118 (2019). doi:10.1101/gr.239277.118

37. Chaudhuri, J. et al. Transcription-targeted DNA deamination by the AID antibody diversification enzyme. Nature (2003). doi:10.1038/nature01574

38. Álvarez-Prado, Á. F. et al. A broad atlas of somatic hypermutation allows prediction of activation-induced deaminase targets. J. Exp. Med. (2018). doi:10.1084/jem.20171738

39. Rogozin, I. B. & Kolchanov, N. A. Somatic hypermutagenesis in immunoglobulin genes. II. Influence of neighbouring base sequences on mutagenesis. BBA – Gene Struct. Expr. (1992). doi:10.1016/0167-4781(92)90134-L

40. Qian, J. et al. B cell super-enhancers and regulatory clusters recruit AID tumorigenic activity. Cell (2014). doi:10.1016/j.cell.2014.11.013

41. Meng, F. L. et al. Convergent transcription at intragenic super-enhancers targets AID-initiated genomic instability. Cell (2014). doi:10.1016/j.cell.2014.11.014

42. Morgan, H. D., Dean, W., Coker, H. A., Reik, W. & Petersen-Mahrt, S. K. Activation-induced cytidine deaminase deaminates 5-methylcytosine in DNA and is expressed in pluripotent tissues: Implications for epigenetic reprogramming. J. Biol. Chem. (2004). doi:10.1074/jbc.M407695200

43. Lio, C. W. J. et al. TET enzymes augment activation-induced deaminase (AID) expression via 5-hydroxymethylcytosine modifications at the Aicda superenhancer. Sci. Immunol. (2019). doi:10.1126/sciimmunol.aau7523

44. Cantaert, T. et al. Activation-Induced Cytidine Deaminase Expression in Human B Cell Precursors Is Essential for Central B Cell Tolerance. Immunity (2015). doi:10.1016/j.immuni.2015.10.002

45. Kuraoka, M. et al. Activation-induced cytidine deaminase mediates central tolerance in B cells. Proc. Natl. Acad. Sci. U. S. A. (2011). doi:10.1073/pnas.1102571108

46. Yang, K. et al. Activation-Induced Cytidine Deaminase Expression and Activity in the Absence of Germinal Centers: Insights into Hyper-IgM Syndrome. J. Immunol. (2009). doi:10.4049/jimmunol.0901548

47. Neri, F. et al. Intragenic DNA methylation prevents spurious transcription initiation. Nature (2017). doi:10.1038/nature21373

48. Ise, W. et al. The transcription factor BATF controls the global regulators of class-switch recombination in both B cells and T cells. Nat. Immunol. (2011). doi:10.1038/ni.2037

49. Betz, B. C. et al. Batf coordinates multiple aspects of B and T cell function required for normal antibody responses. The Journal of experimental medicine (2010). doi:10.1084/jem.20091548

50. Gekas, C. et al. Mef2C is a lineage-restricted target of Scl/Tal1 and regulates megakaryopoiesis and B-cell homeostasis. Blood 113, 3461–3471 (2009).

51. Herglotz, J. et al. Essential control of early B-cell development by Mef2 transcription factors. Blood (2016). doi:10.1182/blood-2015-04-643270

52. Ryan, R. J. H. et al. Detection of enhancer-associated rearrangements reveals mechanisms of oncogene dysregulation in B-cell lymphoma. Cancer Discov. (2015). doi:10.1158/2159-8290.CD-15-0370

53. Kandasamy, K. et al. NetPath: A public resource of curated signal transduction pathways. Genome Biol. (2010). doi:10.1186/gb-2010-11-1-r3

54. Feingold, E. A. et al. The ENCODE (ENCyclopedia of DNA Elements) Project. Science (2004). doi:10.1126/science.1105136

55. Cahir-McFarland, E. D. et al. Role of NF-kappa B in cell survival and transcription of latent membrane protein 1-expressing or Epstein-Barr virus latency III-infected cells. J. Virol. 78, 4108–19 (2004).

56. Caldwell, R. G., Wilson, J. B., Anderson, S. J. & Longnecker, R. Epstein-Barr virus LMP2A drives B cell development and survival in the absence of normal B cell receptor signals. Immunity (1998). doi:10.1016/S1074-7613(00)80623-8

57. Cen, O. & Longnecker, R. Latent Membrane Protein 2 (LMP2). in 151-180 (Springer, Cham, 2015). doi:10.1007/978-3-319-22834-1_5

58. Mancao, C. & Hammerschmidt, W. Epstein-Barr virus latent membrane protein 2A is a B-cell receptor mimic and essential for B-cell survival. Blood 110, 3715–3721 (2007).

59. Rickinson, A. B. & Kieff, E. D. Epstein-Barr virus and its replication. in Fields Virology (2007). doi:10.1016/j.pce.2015.01.007

60. Luo, W., Weisel, F. & Shlomchik, M. J. B Cell Receptor and CD40 Signaling Are Rewired for Synergistic Induction of the c-Myc Transcription Factor in Germinal Center B Cells. Immunity (2018). doi:10.1016/j.immuni.2018.01.008

61. Ma, Y. et al. CRISPR/Cas9 Screens Reveal Epstein-Barr Virus-Transformed B Cell Host Dependency Factors. Cell Host Microbe 21, 580–591.e7 (2017).

62. Li, P. et al. BATF-JUN is critical for IRF4-mediated transcription in T cells. Nature (2012). doi:10.1038/nature11530

63. Ochiai, K. et al. Transcriptional Regulation of Germinal Center B and Plasma Cell Fates by Dynamical Control of IRF4. Immunity (2013). doi:10.1016/j.immuni.2013.04.009

64. Mao, C. et al. T Cell-Independent Somatic Hypermutation in Murine B Cells with an Immature Phenotype. Immunity (2004). doi:10.1016/S1074-7613(04)00019-6

65. Cappione, A. et al. Germinal center exclusion of autoreactive B cells is defective in human systemic lupus erythematosus. J. Clin. Invest. (2005). doi:10.1172/JCI24179

66. Richardson, C. et al. Molecular Basis of 9G4 B Cell Autoreactivity in Human Systemic Lupus Erythematosus. J. Immunol. (2013). doi:10.4049/jimmunol.1202263

67. Moura, R. A. et al. 9G4 expression on autoantibodies to citrullinated peptides in patients with early inflammatory arthritis and established rheumatoid arthritis. Ann. Rheum. Dis. (2012). doi:10.1136/annrheumdis-2011-201231.15

68. Cambridge, G. et al. Expression of the inherently autoreactive idiotope 9G4 on autoantibodies to citrullinated peptides and on rheumatoid factors in patients with early and established rheumatoid arthritis. PLoS One (2014). doi:10.1371/journal.pone.0107513

69. Meyers, G. et al. Activation-induced cytidine deaminase (AID) is required for B-cell tolerance in humans. Proc. Natl. Acad. Sci. (2011). doi:10.1073/pnas.1102600108

70. Goodnow, C. C. Balancing immunity and tolerance: deleting and tuning lymphocyte repertoires. Proc. Natl. Acad. Sci. (1996). doi:10.1073/pnas.93.6.2264

71. Meffre, E., Casellas, R. & Nussenzweig, M. C. Antibody regulation of B cell development. Nat. Immunol. (2000). doi:10.1038/80816

72. Nemazee, D. Receptor editing in lymphocyte development and central tolerance. Nature Reviews Immunology (2006). doi:10.1038/nri1939

73. Cantaert, T. et al. Activation-Induced Cytidine Deaminase Expression in Human B Cell Precursors Is Essential for Central B Cell Tolerance. Immunity (2015). doi:10.1016/j.immuni.2015.10.002

74. Quartier, P. et al. Clinical, immunologic and genetic analysis of 29 patients with autosomal recessive hyper-IgM syndrome due to Activation-Induced Cytidine Deaminase deficiency. Clin. Immunol. (2004). doi:10.1016/j.clim.2003.10.007

75. Roco, J. A. et al. Class-Switch Recombination Occurs Infrequently in Germinal Centers. Immunity 0, (2019).

76. Seidel, M. G. et al. The European Society for Immunodeficiencies (ESID) Registry Working Definitions for the Clinical Diagnosis of Inborn Errors of Immunity. J. Allergy Clin. Immunol. Pract. (2019). doi:10.1016/j.jaip.2019.02.004

77. Li, H. & Durbin, R. Fast and accurate long-read alignment with Burrows-Wheeler transform. Bioinformatics (2010). doi:10.1093/bioinformatics/btp698

78. Liu, H. et al. Systematic identification and annotation of human methylation marks based on bisulfite sequencing methylomes reveals distinct roles of cell type-specific hypomethylation in the regulation of cell identity genes. Nucleic Acids Res. (2016). doi:10.1093/nar/gkv1332

79. Stunnenberg, H. G. et al. The International Human Epigenome Consortium: A Blueprint for Scientific Collaboration and Discovery. Cell (2016). doi:10.1016/j.cell.2016.11.007

80. Li, H. et al. The Sequence Alignment/Map format and SAMtools. Bioinformatics (2009). doi:10.1093/bioinformatics/btp352

81. Broad Institute. Picard Toolkit. http://broadinstitute.github.io/picard/ (2018).

82. Zhang, Y. et al. Model-based analysis of ChIP-Seq (MACS). Genome Biol 9, R137 (2008).

83. Whyte, W. A. et al. Master transcription factors and mediator establish super-enhancers at key cell identity genes. Cell (2013). doi:10.1016/j.cell.2013.03.035

84. Pageaud, Y., Plass, C. & Assenov, Y. Enrichment analysis with EpiAnnotator. Bioinformatics (2018). doi:10.1093/bioinformatics/bty007

85. McLean, C. Y. et al. GREAT improves functional interpretation of cis-regulatory regions. Nat. Biotechnol. (2010). doi:10.1038/nbt.1630

